# No association between plant mating system and geographic range overlap

**DOI:** 10.1101/016261

**Authors:** Dena Grossenbacher, Ryan Briscoe Runquist, Emma E. Goldberg, Yaniv Brandvain

## Abstract

**Premise of the Study:** Automatic self-fertilization may influence the geography of speciation, promote reproductive isolation between incipient species, and lead to ecological differentiation. As such, selfing taxa are predicted to co-occur more often with their closest relatives than are outcrossing taxa. Despite suggestions that this pattern may be general, the extent to which mating system influences range overlap in close relatives has not been tested formally across a diverse group of plant species pairs.

**Methods:** We test for a difference in range overlap between species pairs where zero, one, or both species are selfers, using data from 98 sister species pairs in 20 genera across 15 flowering plant families. We also use divergence time estimates from time-calibrated phylogenies to ask how range overlap changes with divergence time and whether this effect depends on mating system.

**Key Results:** We find no evidence that automatic self-fertilization influences range overlap of closely related plant species. Sister pairs with more recent divergence times had modestly greater range overlap, but this effect did not depend on mating system.

**Conclusions:** The absence of a strong influence of mating system on range overlap suggests that mating system plays a minor or inconsistent role compared to many other mechanisms potentially influencing the co-occurrence of close relatives.

## INTRODUCTION

From the initiation to the completion of speciation and beyond, mating system can dramatically influence the potential for gene exchange, competition for pollinators, and ecological differentiation (Antonovics, 1968; Levin, 1972; Jain, 1976; Fishman and Wyatt, 1999; Brandvain and Haig, 2005; Martin and Willis, 2007; Smith and Rausher, 2007; Briscoe Runquist and Moeller, 2013). Self-fertilization can alter the geographic mode of speciation, the extent of reproductive isolation between incipient species, and the subsequent sympatric persistence of sister species. Consequently, autonomous self-fertilization may influence the extent of co-occurrence of closely related species, and mating system could serve as a model for understanding the role of functional traits in speciation. Although case studies and evolutionary theory both suggest that selfing can allow closely related plant species to co-occur (Antonovics, 1968; Whalen, 1978; Levin, 1985; Fishman and Wyatt, 1999; Martin and Willis, 2007; Smith and Rausher, 2007; Levin, 2010; Matallana et al., 2010; Grossenbacher and Whittall, 2011; Briscoe Runquist and Moeller, 2013; Vallejo-Marin et al., 2014), the generality of the hypothesis that range overlap is greater in pairs of species in which one or both is selfing has not been tested formally across a diverse group of plant species pairs.

Mating system can influence patterns of range overlap by *(i) its influence on the geographic context in which species arise, (ii) preventing species’ fusion upon secondary contact by promoting reproductive isolation, and (iii) enabling co-existence by ecological differentiation.* We discuss these in turn.

### Geography of speciation

There are at least three plausible scenarios under which autonomous self-fertilization would influence the geographic mode of speciation and consequently the range overlap of recently diverged selfing-outcrossing and selfing-selfing sister pairs. In the first scenario, selfing species arise following long distance dispersal. Because autonomous selfing allows rare migrants to colonize and establish (Baker’s law; Baker, 1955), a sole migrant experiencing a long-distance dispersal event can give rise to an entire selfing species that is allopatric from its closest relative. Baker’s law thus suggests a filter by which mating system may influence speciation. This is thought to be the case in *Capsella* (Foxe et al., 2009; Guo et al., 2009; but see Brandvain et al., 2013) and in the sea star *Cryptasterina hystera* (Puritz et al., 2009). In the second scenario, selfing may be favored by selection as a means to provide reproductive assurance in marginal habitats (Lloyd, 1992; Schoen, 1996) just outside of the range of an outcrossing relative. This scenario could result in peripatric speciation, as may be the case in *Clarkia* (Lewis and Lewis, 1955; Moeller and Geber, 2005). In the third scenario, selfing may evolve in a population adapted to a novel habitat directly adjacent to an outcrossing population,serving as a mechanism to shield locally adaptive genomes from maladaptive introgression (Levin, 2010). This may be the case in several grass species (Antonovics, 1968), *Mimulus* (Ferris et al., 2014), and *Layia* (Baldwin, 2005). In all three scenarios, selfing may either evolve concurrently with colonization or ecological adaptation (producing a selfing-outcrossing sister pair), or it may already be the mating system of the parental species (producing a selfing-selfing sister pair). In the two latter scenarios, selfing populations and species arise geographically nearby their close relatives, so subsequent range shifts or range expansion in selfers (e.g., Grossenbacher et al., 2015) may lead to range overlap and increased amounts of secondary contact. The relationship between mating system and range overlap may thus depend on the time elapsed since speciation.

### Reproductive isolation

Autonomous self-fertilization limits gene flow, promotes reproductive isolation, and maintains the distinctness of recently diverged lineages in several ways, each of which could facilitate co-existence. Perhaps most importantly, the transition toward selfing is generally associated with reductions in pollinator attraction traits and reduced visitation by pollinators (reviewed in Sicard and Lenhard, 2011), decreasing opportunities for heterospecific pollen movement between predominantly selfing and predominantly outcrossing taxa (e.g., Fishman and Wyatt, 1999; Martin and Willis, 2007). In fact, in some cases selfing may evolve or be enhanced following secondary contact as a means to prevent the formation of maladaptive hybrids (reinforcement), as is likely in *Clarkia xantiana* (Briscoe Runquist and Moeller, 2013). In cases where pollen transfer does occur, pollen-pistil incompatibilities and abnormal seed development may pleiotropically follow the evolution of selfing (Brandvain and Haig, 2005; Koelling et al., 2011), further reducing the chance of successful hybridization. Together, these barriers could lead to near-complete reproductive isolation between selfing and outcrossing taxa, preventing their fusion.

### Ecological coexistence

In addition to potentially promoting reproductive isolation, selfing can facilitate the ecological coexistence of closely related species by reducing pollinator competition. Many studies across angiosperms document that competition for pollinator services can have massive impacts on fitness, population establishment, and persistence (e.g., Waser, 1978; Fishman and Wyatt, 1999; Brown et al., 2002; Bell et al., 2005; Briscoe Runquist, 2012; Grossenbacher and Stanton, 2014). Predominant selfing may eliminate pollinator-mediated competition by reducing reliance on pollinators altogether, allowing species to coexist and preventing competitive exclusion following secondary range shifts. Experimental field transplants have demonstrated the potential importance of this mechanism of co-existence. In *Mimulus ringens* (Bell et al., 2005), competition for pollinator services with an invasive species caused reduced conspecific pollen deposition; plants compensated for the reduction in fitness through a facultative increase in autonomous selfing. In the typically bee-pollinated *Arenaria* (Fishman and Wyatt, 1999) and *Ipomoea* (Smith and Rausher, 2007), competitive interference due to heterospecific pollen transfer from congeners generated female fitness costs that favored increased selfing.

### Evidence to date

Many biologically plausible models suggest that selfing and outcrossing species will be likely to co-occur. Numerous compelling case studies support this prediction. For example, in Texas, populations of *Phlox drummondii* showed increased self-compatibility in sympatry with its close relative *P. cuspidata* (Levin, 1985). In Mexico, *Solanum grayi* has dramatically reduced flowers and increased selfing rates when it occurs sympatrically with its close relative *S. lumholtzianum*, a pattern that may exist between other closely related species in this clade (Whalen, 1978; Vallejo-Marin et al., 2014). Similarly, populations of *Arenaria* in the southeastern United States and populations of *Clarkia* in southern California have increased selfing rates in sympatry with closely related congeners (Fishman and Wyatt, 1999; Briscoe Runquist and Moeller, 2013). In the genus *Mimulus*, sister species that include one selfing species (selfing-outcrossing sister species) are more likely to occur sympatrically than are outcrossing-outcrossing sister species given similar amounts of divergence time (Grossenbacher and Whittall, 2011). Finally, among the Bromeliads in southeastern Brazil, self-compatible species co-occur with significantly more con-familials than do self-incompatible species (Matallana et al., 2010).

Although these case studies suggest that selfing facilitates co-occurrence of closely related species, the influence of mating system on range overlap has not been tested at a scale larger than focal genera. Here, we test the hypothesis that selfing facilitates co-occurrence of close relatives by asking whether, across many pairs of sister species, co-occurrence is greater or lesser for pairs that contain a selfer. We then use divergence time estimates from time-calibrated phylogenies to explore whether the extent of co-occurrence changes with divergence time, reflecting the extent of post-speciational range shifts, and whether this effect depends on mating system. Surprisingly, our results do not support the anecdotal relationship between mating system and range overlap. We suggest that although in some instances mating system has a major influence on range overlap, the overall effect is weak, inconsistent, and does not scale up from microevolutionary process to macroevolutionary and macroecological pattern.

## MATERIALS AND METHODS

We identified taxa with a published, species-level phylogeny containing at least one predominantly selfing or functionally selfing species and one predominantly outcrossing species, and with DNA sequence data for at least 50% of the species within the clade available on GenBank (http://www.ncbi.nlm.nih.gov/genbank/) to be used for constructing time-calibrated phylogenies. After removing *Leavenworthia*, a small North American genus in which our phylogenetic model did not converge, we had 20 clades from 15 families whose combined native distributions spanned every continent except Antarctica (Appendix S1, see Supplemental Data with the online version of this article). On average, clades contained 35 ±7 (±1SE) extant species, 80 ±4.6 percent of which were included in our phylogenies. These time-calibrated, species level phylogenies across a diverse set of plant taxa allow us to test whether mating system influences species’ co-occurrence, while controlling for shared evolutionary history.

For the analyses described below, all data and R scripts are available from the Dryad Digital Repository (doi:10.5061/dryad.hv117; Grossenbacher et al., 2015). We previously described our data set, phylogeny estimation, and data for species’ traits in a separate analysis of the question of how mating system influences range size (Grossenbacher et al., 2015).

### Identifying sister species pairs

We generated time-calibrated phylogenies for all 20 genera or generic sections using publicly available sequence data. We reconstructed phylogenies because most previously published phylogenies were not time calibrated and consisted of only a single topology or consensus tree, making it difficult to incorporate uncertainty into our analysis. Prior to estimating the phylogenies, for each clade separately, we downloaded sequences for the nuclear ribosomal internal transcribed spacer locus (nrITS) for species within the clade from GenBank and aligned them using the MUSCLE package in R, version 3.8.31-4 (Edgar, 2004). We simultaneously estimated the phylogenetic relationships and the absolute divergence times among species in a Bayesian framework in BEAST version 1.6.2 (Drummond et al., 2012). To estimate absolute divergence times, we used the mean and range of substitution rate for herbaceous and woody plants at the nrITS locus (Kay et al., 2006), because fossils are not known for any of the clades in the analysis. As in Grossenbacher et al. (2015), we set the substitution rate to a normally distributed prior for herbaceous lineages with mean of 4.13 × 10^-9^ subs/site/yr and standard deviation of 1.81 × 10^-9^, and for woody lineages with mean of 2.15 × 10^-9^ subs/site/yr and standard deviation of 1.85 × 10^-9^.

To accommodate heterogeneity in the molecular evolutionary rate among branches, we used an uncorrelated log-normal relaxed clock model. The prior model on branch lengths was set to a Yule process of speciation. The prior model on substitutions and the number of MCMC generations varied by clade (see Appendix S2, see Supplemental Data with the online version of this article). Posterior samples of parameter values were summarized and assessed for convergence and mixing using Tracer v. 1.5 (Rambaut et al., 2014). After removing *Leavenworthia* (for which the MCMC did not converge, and which we excluded from all analyses), all MCMCs for phylogenies of our 20 clades had minimum estimated sum of squares (ESS) for the posterior >1100, and minimum ESS across all other parameters >600 (Appendix S2).

We identified sister species in a subset of 9000 trees from the posterior distribution for each clade. For each sister species pair, we recorded the average divergence time and the posterior probability of that pair as the proportion of trees that contained that pair, providing a measure of phylogenetic uncertainty. Since our phylogenies sampled, on average, only 80% of extant taxa, these sister pairs may not represent “true” extant sisters, but they are recently diverged groups representing independent evolutionary replicates. For all ensuing analyses, we used the identified sister pairs that had the highest posterior probabilities and did not duplicate species already in the dataset, to avoid pseudoreplication.

### Estimating mating system, ploidy, and lifespan

We collated 54 studies that described the mating systems of species from the 20 genera or generic sections identified above. Most published studies classified species as predominantly selfing, variable mating, or predominantly outcrossing. As in Grossenbacher et al. (2015), we classified species as variable mating when outcrossing rates within an individual or population were between 0.2 and 0.8, or when there was extensive among-population variation in outcrossing rates. An exception to this classification scheme were species in *Oenothera* sect. *oenothera*, which were classified as either sexually reproducing or functionally asexual, due to a permanent translocation whereby plants self-fertilize but do not undergo segregation and recombination (Johnson et al., 2009). Sexual *Oenothera* sect. *oenothera* species are partially or wholly self-incompatible, and they are assumed to be outcrossing relative to the asexual species. Methods for mating system classification varied among clades because different traits are more reliable indicators of mating system in different taxa; within clades methods were generally consistent (Appendix S3, see Supplemental Data with the online version of this article). To extend our data set, we occasionally classified taxa that were missing from the primary studies using the same traits and metrics as those used for other species within that clade (Appendix S3). We then assigned previously identified sister pairs to one of three mating system categories: outcrosser-outcrosser, selfer-outcrosser, or selfer-selfer. Pairs that included variable mating species were excluded from this analysis.

Mating system may coevolve and be correlated with traits such as polyploidy (Stebbins, 1950; Barringer, 2007, Robertson et al., 2011) and lifespan (Barrett et al., 1996). To ensure that these traits did not drive or obscure a relationship between mating system shifts and co-occurrence, we gathered published information on ploidy and lifespan when possible. For ploidy, we recorded chromosome counts and classified each species (relative to the base ploidy reported for each genus in the literature) as diploid, polyploid, or mixed when both diploid and polyploid individuals were known. Species’ lifespans were classified as annual, perennial, or mixed when both annual and perennial individuals were known

### Estimating co-occurrence / geographic range overlap

We downloaded all known species occurrence records for the clades from the Global Biodiversity Information Facility (http://www.gbif.org) and filtered for quality by excluding records with coordinate accuracy <100 km, coordinates failing to match the locality description, and taxonomic misidentifications (verified by the authors and taxonomic specialists of each clade). We checked species’ epithets against the most recently published taxonomies and corrected synonyms and spelling errors. We included only coordinates from the native range of species. Coordinates outside the native species range were identified using published monographs and online databases that report native and invaded ranges (e.g., GRIN database, http://www.ars-grin.gov/).

We used the filtered occurrence data to estimate the degree of co-occurrence using a grid approach. We divided the world into a series of rectangular cells by grid lines that follow degree longitude and latitude using the “raster” R package version 2.3-0 (Hijmans et al., 2011). We calculated co-occurrence as the summed area of grid cells occupied by both species, divided by the summed area of occupied grid cells for the smaller ranged species. Thus, co-occurrence ranges between 0 (no range overlap) and 1 (the smaller-ranged species is found only within the range of the larger-ranged species) (Barraclough and Vogler, 2000; Fitzpatrick and Turelli, 2006). In order to assess whether the ensuing analyses were sensitive to the spatial scale at which co-occurrence is estimated, co-occurrence was calculated across a range of cell sizes, 0.05, 0.1, 0.5 and 1 decimal degrees, representing grid cells of roughly 25, 100, 2500, and 10000 km^2_2_^ respectively (exact value varies by latitude), again as Grossenbacher et al. (2015).

### Analyses

To explore whether divergence time varied by mating system, we used analysis of variance (ANOVA). To meet model assumptions, the response variable (divergence time) was natural log-transformed prior to analysis. The predictor variable (sister pair mating system) was categorical with three states: outcrosser-outcrosser, selfer-outcrosser, and selfer-selfer. To incorporate phylogenetic uncertainty into our analysis, and all subsequent models, we included a weighting factor for each sister pair that was equal to the posterior probability of the sister pair (the proportion of phylogenetic trees that contained a given sister pair).

To test whether the mating system of species pairs influences co-occurrence, we used beta regression models in the ‘betareg’ package in R (Cribari-Neto and Zeileis, 2009). Beta regression provides a flexible model for continuous response variables defined on the interval (0,1) that display both heteroscedasticity and skewness, e.g., proportional data with many values close to zero. The response variable (co-occurrence) was transformed prior to analysis, using a standard transformation on co-occurrence values (y(n-1) + 0.5/n where n is the sample size, Smithson and Verkuilen, 2006) because in some cases co-occurrence assumed values of 0 and 1. The predictor variable (sister pair mating system) was categorical with three states as described above. We fit this model using maximum likelihood with a bias correction to determine confidence intervals of the estimated coefficients. We used partial Wald tests to compare among the three mating system categories.

To determine whether time since divergence influences co-occurrence, we used beta regression as in the model described above, where the response variable was transformed co-occurrence and the predictor variable was log divergence time. To determine whether the relationship between co-occurrence and divergence time varied by sister pair mating system, we added two additional predictors to this model: sister pair mating system and its interaction with divergence time.

To examine whether our results were robust to the spatial scale at which co-occurrence was determined, we performed all analyses four times using the four grid cell sizes described above. We also ran all analyses including only sister pairs that did not differ in ploidy and lifespan to ensure that our results were not driven by these potentially correlated traits. Finally, to explore the possibility that certain clades were heavily influencing overall results, we ran all models described above while sequentially dropping individual clades (N=20). We report cases where dropping a single clade altered the significance of any model effects.

## RESULTS

We identified 98 sister species pairs from the phylogenetic analysis across 20 genera and generic sections. Of these pairs, 52 were outcrossing-outcrossing, 30 were selfing-outcrossing, and 16 were selfing-selfing.

Divergence time varied across mating system categories, with outcrosser-outcrosser sister species roughly two times older, on average, than selfer-outcrosser sister species (Fig. 1; overall ANOVA, *F*=4.962_2,95_, *P*=0.009; Tukey LSM difference test, outcrosser-outcrosser – selfer-outcrosser *P* = 0.011, outcrosser-outcrosser – selfer-selfer *P* = 0.482, selfer-outcrosser – selfer-selfer *P* = 0.506); consistent with the notion of selfing as an “evolutionary dead end” (see Stebbins 1957, Takebayashi and Morrell, 2001, Igic and Busch, 2013). There was large variation in co-occurrence for all ‘sister pair mating system’ categories, especially for young sister pairs.

**Figure 1.**
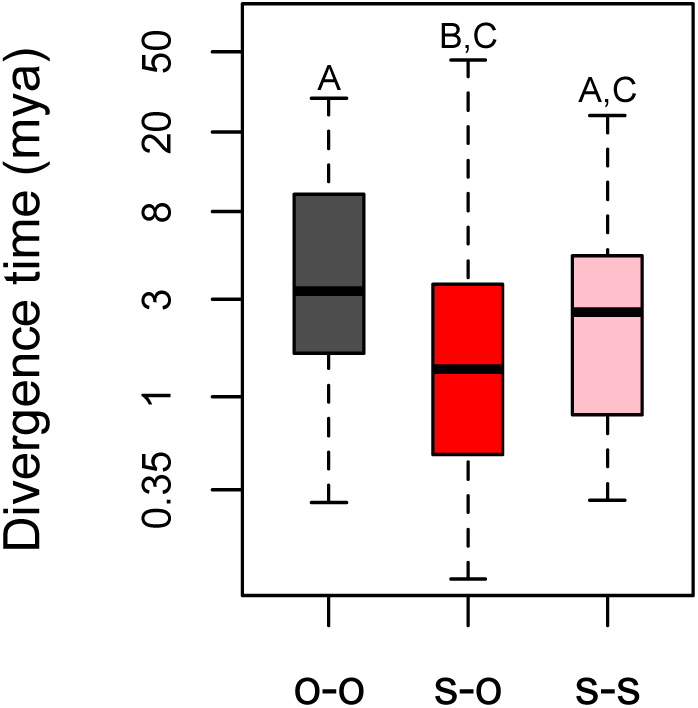
Box plots of sister pair divergence times by mating system category: outcrossing-outcrossing (o-o, dark gray), selfing-outcrossing (s-o, red), selfing-selfing (s-s, pink). Letters represent *a posteriori* Tukey groupings; see text for ANOVA summary. Divergence time axis is natural logarithmic scale (back-transformed).

Patterns of co-occurrence between sister species were not strongly influenced by their mating systems. The distribution of co-occurrences between sister species ranged from zero to one, and it was considerably skewed toward zero across all mating system categories (Fig. 2). Only at the finest spatial scale did mating systems of sister pairs explain even a marginally significant proportion of the variation in co-occurrence— selfing-selfing sisters had, on average, about two times greater co-occurrence than outcrossing-outcrossing sisters (*P* = 0.065; Table 1; Fig. 2). However, this result is largely driven by a single clade, *Medicago*, which contained 5 selfer-selfer pairs. When *Medicago* was dropped from the analysis, the effect of selfing-selfing sisters on co-occurrence disappeared (*P* = 0.504). These results were not qualitatively different after excluding sister pairs that differed in ploidy and life span (not presented).

**Figure 2.**
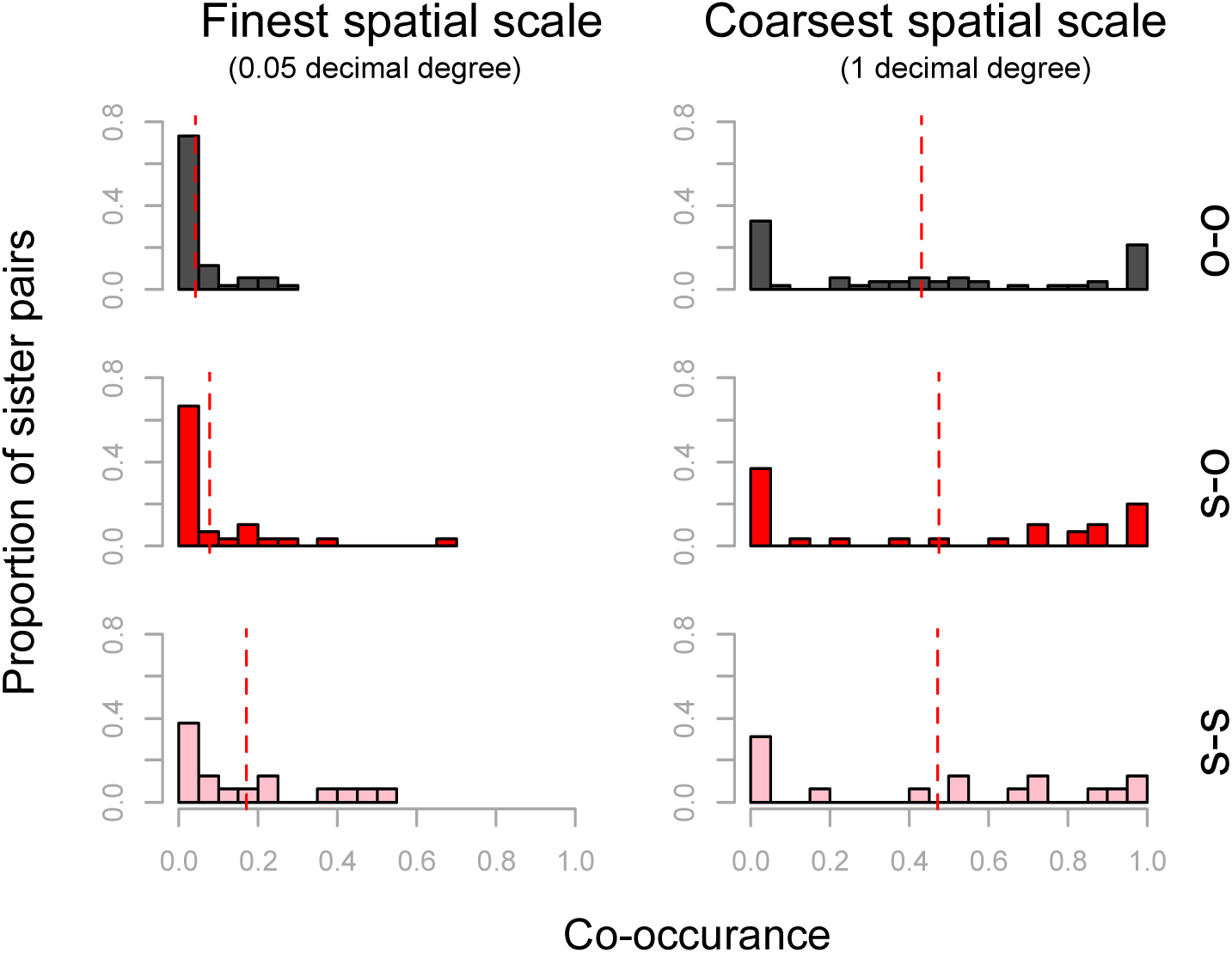
Histograms of sister pair co-occurrence by mating system category: outcrossing-outcrossing (o-o, dark gray), selfing-outcrossing (s-o, red), selfing-selfing (s-s, pink). Dashed vertical lines indicate mean co-occurrence. See Table 1 for statistical results.

**Table 1.** Results of beta regression models analyzing the effect of ‘sister pair mating system’ on co-occurrence, estimated at four spatial scales (**A-D**).

|  | Coefficient<br>(s.e.) | Wald's<br>*z-value* | *P* |
| --- | --- | --- | --- |
| **A. 0.05 decimal degree (~25 km<sup>2</sup>)** |  |  |  |
| Intercept | -2.371 (0.253) | -9.377 | <0.001 |
| selfer-outcrosser | 0.031 (0.315) | 0.098 | 0.922 |
| selfer-selfer | 0.646 (0.350) | 1.844 | 0.065 |
| **B. 0.1 decimal degree (~100 km<sup>2</sup>)** |  |  |  |
| Intercept | -2.003 (0.254) | -7.878 | <0.001 |
| selfer-outcrosser | 0.209 (0.326) | 0.640 | 0.522 |
| selfer-selfer | 0.606 (0.369) | 1.643 | 0.100 |
| **C. 0.5 decimal degree (~2500 km<sup>2</sup>)** |  |  |  |
| Intercept | -0.581 (0.268) | -2.168 | 0.030 |
| selfer-outcrosser | 0.482 (0.413) | 1.168 | 0.243 |
| selfer-selfer | 0.211 (0.462) | 0.457 | 0.648 |
| **D. 1 decimal degree (~10,000 km<sup>2</sup>)** |  |  |  |
| Intercept | -0.071 (0.271) | -0.263 | 0.792 |
| selfer-outcrosser | 0.220 (0.420) | 0.523 | 0.601 |
| selfer-selfer | 0.084 (0.476) | 0.177 | 0.860 |

Note: The categorical coefficient estimates are log-odds ratios, measured as departures from the ‘outcrosser-outcrosser’ category.

Although the distribution of divergence times differed between the three mating system categories (Fig. 1), the relationship between divergence time and range overlap did not obscure the effect of mating system on co-occurrence. There was a weak trend for co-occurrence to increase with decreased divergence time, but only at the coarsest spatial scale, and even then divergence time explained only a marginally significant proportion of the variation in co-occurrence (*P* = 0.064; Table 2; Fig. 3). Additionally, when including divergence time in the model with mating system, mating system is not significant (P > 0.286 in all comparisons, Appendix S4, see Supplemental Data with the online version of this article), and the interaction between divergence time and mating system did not influence co-occurrence at any spatial scale (P > 0.295 in all comparisons, Appendix S4). Excluding sister pairs that differed in ploidy and life span did not qualitatively alter these results (not presented). Together, our results do not support the hypothesis that mating systems consistently influence range overlap.

**Figure 3.**
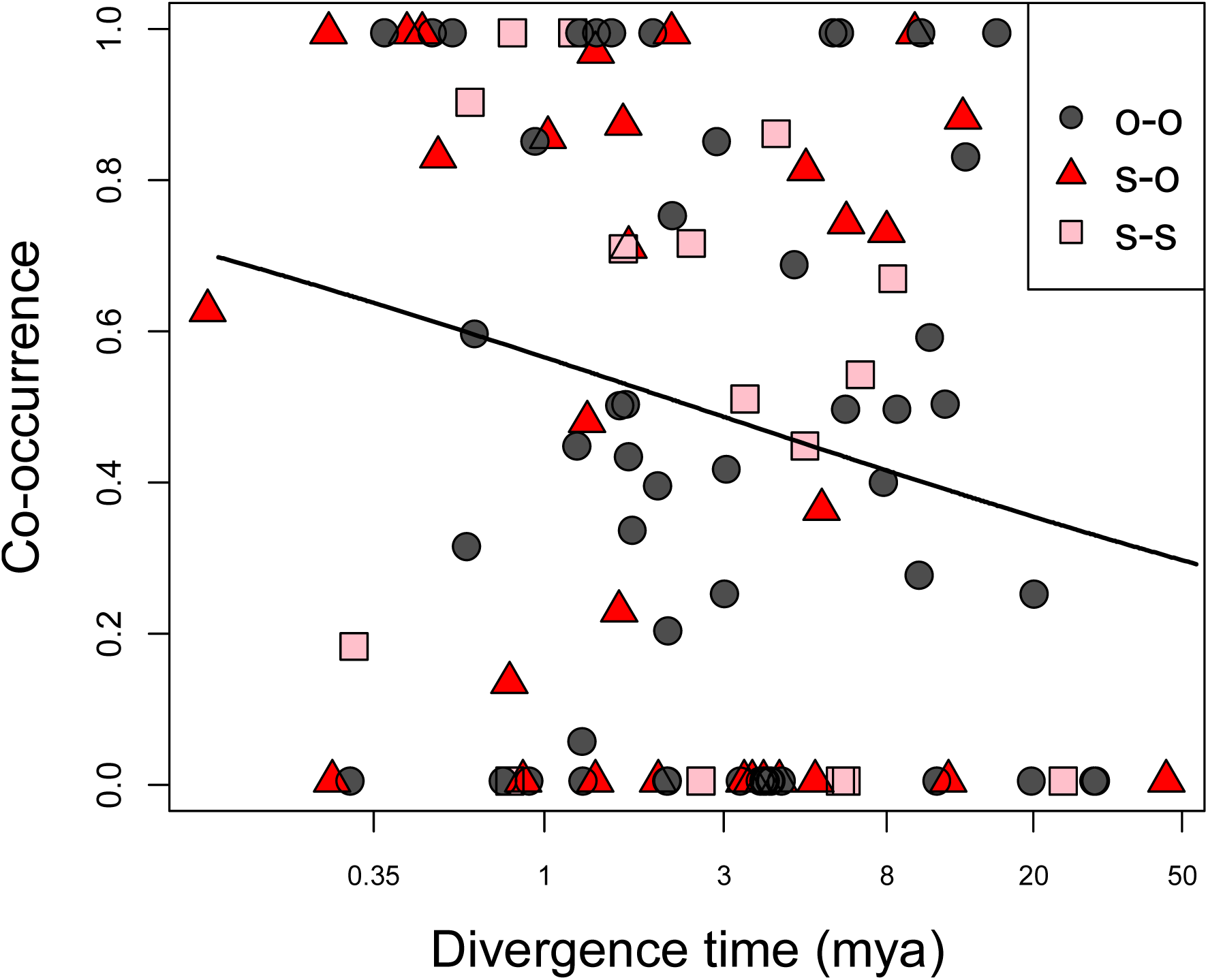
Co-occurrence at the coarsest spatial scale (1 decimal degree) by divergence time for 98 sister species across 20 clades. The line segment represents the predicted slope from beta regression. Divergence time axis is natural logarithmic scale (back-transformed). See Table 2 for statistical results.

**Table 2.** Results of beta regression models analyzing the effect of divergence time on co-occurrence, estimated at four spatial scales (**A-D**).

|  | Coefficient<br>(s.e.) | Wald's<br>*z-value* | *P* |
| --- | --- | --- | --- |
| **A. 0.05 decimal degree (~25 km<sup>2</sup>)** |  |  |  |
| Intercept | -2.118 (0.218) | -9.734 | <0.001 |
| Log divergence time | -0.100 (0.117) | -0.853 | 0.393 |
| **B. 0.1 decimal degree (~100 km<sup>2</sup>)** |  |  |  |
| Intercept | -1.650 (0.212) | -7.800 | <0.001 |
| Log divergence time | -0.181 (0.121) | -1.493 | 0.135 |
| **C. 0.5 decimal degree (~2500 km<sup>2</sup>)** |  |  |  |
| Intercept | -0.166 (0.222) | -0.748 | 0.455 |
| Log divergence time | -0.257 (0.152) | -1.694 | 0.090 |
| **D. 1 decimal degree (~10,000 km<sup>2</sup>)** |  |  |  |
| Intercept | -0.263 (0.224) | 1.173 | 0.241 |
| Log divergence time | -0.287 (0.155) | -1.852 | 0.064 |

## DISCUSSION

### No consistent influence of plant mating system on range overlap

Intuition, theory, and case studies all suggest that autonomous self-fertilization will facilitate range overlap of closely related species (Antonovics, 1968; Whalen, 1978; Levin, 1985; Fishman and Wyatt, 1999; Martin and Willis, 2007; Smith and Rausher, 2007; Levin, 2010; Matallana et al., 2010; Grossenbacher and Whittall, 2011; Briscoe Runquist and Moeller, 2013; Vallejo-Marin et al., 2014). The numerous mechanisms potentially promoting increased co-occurrence between selfers and their close relatives are diverse. First, if selfing species originate cladogenetically (as is the case for the evolution of self-compatibility, see Goldberg and Igic, 2012) in peri- or parapatry, a minor range shift after speciation could promote early secondary range overlap. Next, upon secondary contact, enhanced reproductive isolation conferred by selfing may prevent fusion. Subsequently, reduced competition for pollinators by selfing plants, perhaps enhanced by character displacement or reinforcement selection, may minimize competitive exclusion.

Yet, in an analysis across 20 genera and generic sections, we uncovered no consistent signal of mating system influencing either the geographic mode of speciation or the amount of range overlap. Why then have these plausible mechanisms not combined to generate a strong influence of mating system on range overlap? We reconcile our findings with prior expectations by considering alternative explanations, interpretations, and implications.

### Geography of speciation and mating system

We detected an overall effect of divergence time on range overlap (an age-range correlation), with more range overlap between recently diverged sister species than between distantly diverged sisters. However, the total variation in overlap explained by divergence time is minimal, and the degree of range overlap among recently diverged sister pairs is highly variable, ranging from complete allopatry to complete sympatry at the coarsest spatial scale. Although this pattern of a negative age-range correlation has been widely interpreted as evidence of a ‘sympatric’ mode of speciation (e.g., Barraclough and Vogler, 2000; Fitzpatrick and Turelli, 2006; Anacker and Strauss, 2014), we caution that it can also be generated by the geographic context of extinction. For example, if extinction is more likely for sister species with sympatric or parapatric ranges (e.g., due to competition), older sisters would tend to be the allopatric survivors, producing the negative age-range correlation we observe.

We found no evidence that autonomous self-fertilization affected the predominant geographic mode of speciation-- that is, the relationship between divergence time of sister species and range overlap did not vary with mating system. More specifically, our findings did not support the predominance of any particular geographic mode of speciation associated with the transition to selfing. This implies that allopatric, peri- or parapatric, and sympatric speciation may all occur for selfing-outcrossing sister pairs. Therefore, the evolutionary transition to selfing may have a more complex influence on the geography of speciation than is generally appreciated. In many verbal and quantitative models of the origin of selfing species (e.g., Grant, 1971; Jain, 1976; Lloyd, 1992; Schoen, 1996; Moeller and Geber, 2005), selfers are thought to arise via parapatric (or peripatric) speciation in extreme environments at or beyond the margins of the range of an outcrossing relative. In these scenarios, slight perturbations in the range could generate high levels of range overlap, but this result is inconsistent with our data. In an opposing model, selfers are more likely to arise following long-distance dispersal because of their greater capacity to reproduce when even a sole migrant lands in a new location (Baker, 1955). If common, then selfers may have less present day range overlap with their closest relative simply because of the large (initial) spatial isolation from their closest relatives. This, however, is also not supported by our data.

Ecological differences between selfing and outcrossing species could reconcile our results with prevailing wisdom of the geography of speciation in selfers. Selfing species often exhibit a suite of traits, such as early flowering and drought resistance, that reflect niche differentiation from outcrossers that is consistent but not tied to the mating system *per se* (Guerrant, 1989; Snell and Aarssen, 2005; Sicard and Lenhard, 2011). Thus, if a shift in mating system is associated with local adaptation, environmental filtering may prevent sympatry even during post-speciation range shifts, with selfers remaining in locations that lack pollinators altogether, or locations with harsh environments (e.g., thin rapidly drying soils) that favor rapid growth. Consistent with this explanation, at fine spatial scales we found modestly greater co-occurrence for selfing-selfing sister pairs, particularly in the genus *Medicago*.

### Broad findings versus exceptional case studies

Studies in several taxa demonstrate that upon secondary contact, selfing can be favored as either a mechanism to prevent maladaptive hybridization (reinforcement) or to avoid competition for pollinators (character displacement) (e.g., Fishman and Wyatt, 1999; Smith and Rausher, 2007; Briscoe Runquist and Moeller, 2013). Why then did we not observe greater range overlap in pairs where one or both species was selfing?

One potential explanation is that case studies researching mating system’s role in species coexistence were not selected at random, but rather, were chosen because they highlighted interesting biological phenomena. In our larger data set, these few cases in which selfing facilitated coexistence would be overwhelmed by the less compelling cases. Alternatively, mating system may play an important role in maintaining species distinctness upon secondary contact, but countervailing forces (e.g., niche convergence in selfing species, see above) could overwhelm this signal.

Taxonomic scale may provide another plausible explanation for the discrepancy between our broad species-level results and system-specific studies. Even if reinforcement or character displacement on mating system is common across angiosperms, its importance might be limited to within-population scales. Accordingly, population-level analyses would find an excess of selfing populations in sympatry with populations of a closely related species. Across the entirety of the species range however, the species would be considered variable mating and excluded from our analysis. This pattern of population-level variation in autonomous selfing rate for sympatric versus allopatric populations is found in many taxa. In the more highly selfing subspecies of *Clarkia xantiana*, *C. x. parviflora*, sympatric populations have smaller flowers with higher selfing rates, probably as a result of reinforcing selection, whereas allopatric populations maintain some ability to receive and export outcrossed pollen (Briscoe Runquist and Moeller, 2013). This is also the case in *Arenaria uniflora*, where there is strong selection for autonomous selfing, and selfing populations only occur in areas of sympatry with the close relative *A. glabra* (Fishman and Wyatt, 1999). This would imply that selfing is a potentially important mechanism underlying coexistence, but that this does not generate a discernable macroevolutionary pattern.

### Many ways to coexist or exclude

A final explanation for the lack of a relationship between range overlap and mating system is that mating system is simply one of myriad potential mechanisms that allow close relatives to co-exist. Of the ∼40% of sister pairs that co-occur in the same grid cell in our fine-scale analysis, habitat differences, flowering time differences, pollinator shifts, and post-pollination incompatibilities could also prevent hybridization or competition. For example, pollinator shifts in outcrossing-outcrossing sister species could facilitate their co-existence. Like selfing, pollinator shifts can influence reproductive isolation, competition, and the geography of speciation (reviewed in Kay and Sargent, 2009). If sympatric outcrossing-outcrossing species pairs are enriched for pollinator shifts, then perhaps both selfing and pollinator shifts may encourage co-existence. Pushing this argument one step further, perhaps sympatric close relatives have diverged in various key traits that allows their co-existence, and therefore a test of any particular trait across a large number of angiosperm species pairs will not uncover a systematic effect.

### Temperate study bias

It is worth noting that most species examined in past case studies have temperate rather than tropical distributions (but see Matallana et al., 2010), and 18 of 20 clades in the present study are temperate. Nonetheless, the two clades containing species with largely tropical distributions included here (*Dalechampia* and *Schiedea*) were consistent with our overall results and did not display any relationship between mating system and co-occurrance. Since biotic interactions are predicted to be stronger in the tropics (Schemske et al., 2009), it will be valuable to test globally the relationship between mating system and co-occurrence as more data become available.

### Conclusion: the influence of mating systems on co-occurrence

Ultimately, we find no evidence that mating system consistently influences the geography of speciation or secondary range overlap. Although mating system has a major effect on sympatry in some case studies, there is no discernable effect across the 20 genera and generic sections examined here. Instead, co-occurrence of close relatives may be influenced by many mechanisms, of which transitions to selfing are only a small part. It is also possible that the evolution of selfing is associated with reproductive assurance during adaptation to marginal or mate-limited habitats and is therefore concomitant with other adaptations that preclude general co-occurrence. Alternatively, selection for selfing in secondary contact may be a population level phenomenon that does not scale up to species-level patterns of co-occurrence. Greater understanding of the evolutionary causes of the transition to selfing is necessary to determine the general influence of mating system on co-occurrence.

## Acknowledgements

The authors thank Boris Igic and April Randle for stimulating discussions. We thank Anne Worley, Barbara Neuffer, Andress Franzke, Jeremiah Busch, and Justen Whittall for expert advice on mating systems and phylogeny construction.

---

**Appendix S1.**
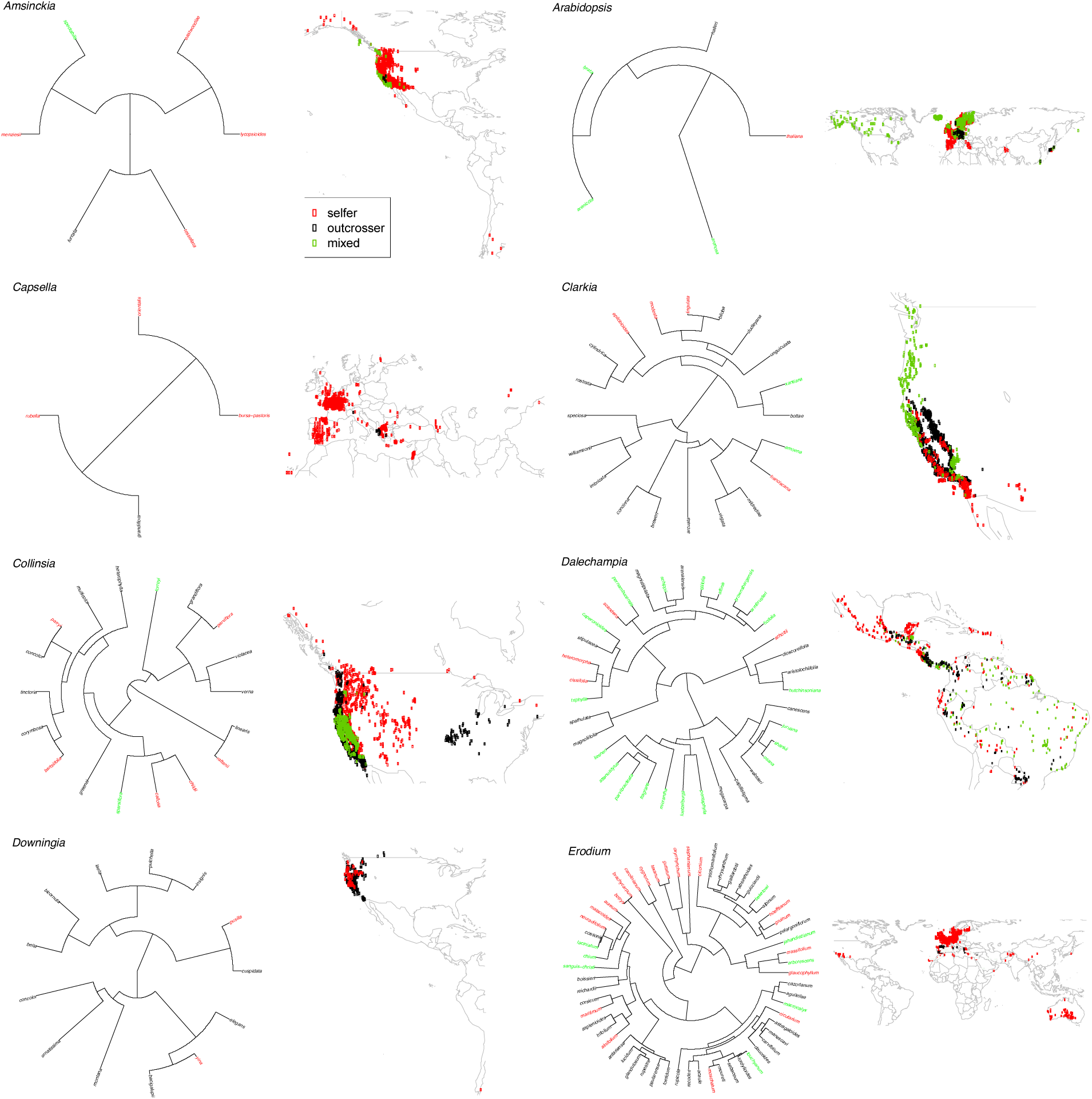
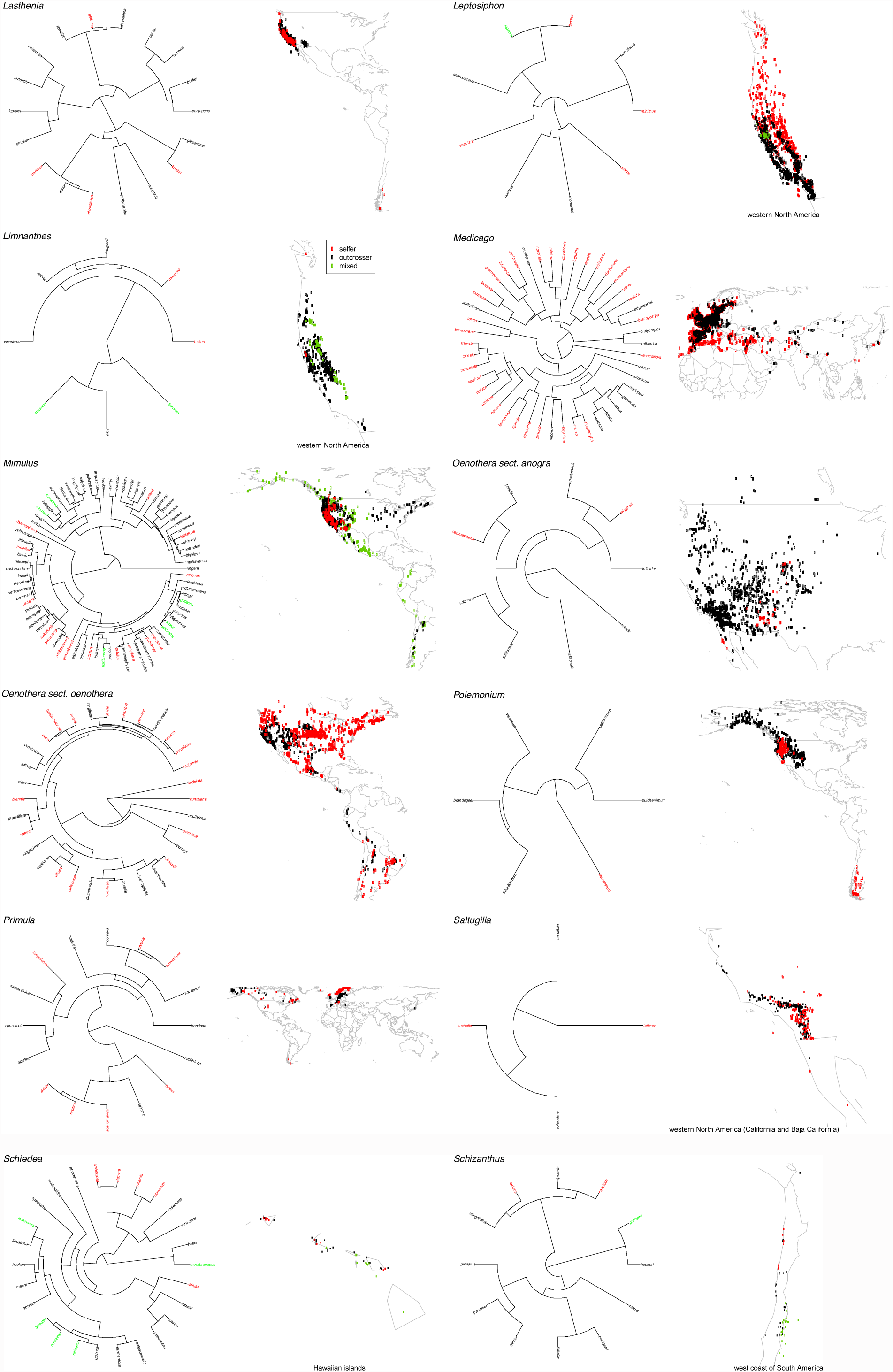
Evolutionary relationships and native distributions of 20 clades. Trees represent bayesian consensus phylogenies with tips colored by mating system (red 641 selfers, black outcrossers, green variable). Geographic distributions represent species’ 642 occurrences, obtained from the global biodiversity information facility (www.gbif.org).

**Appendix S2.**
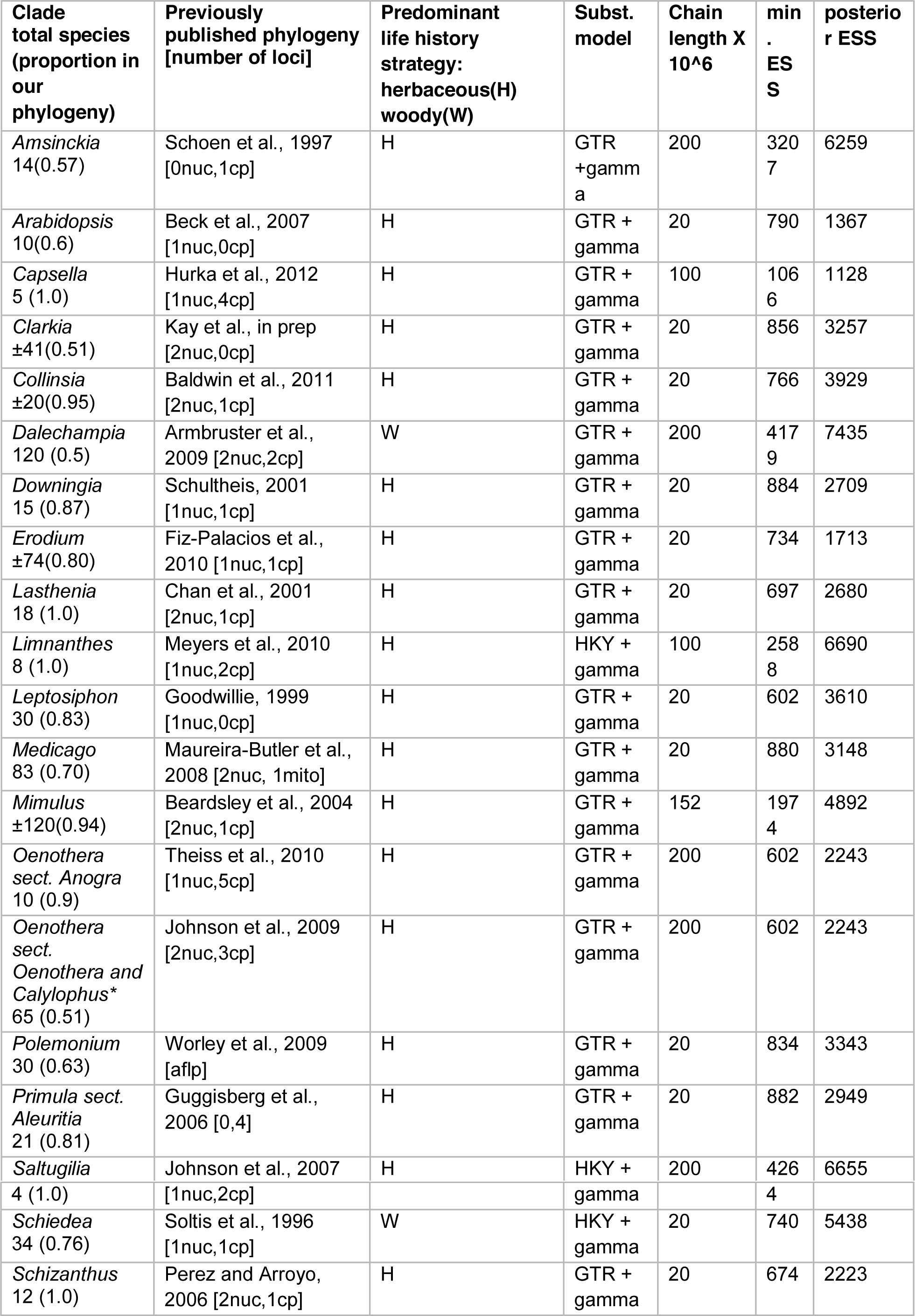
Phylogenetic information for 20 clades included in our study.

**Appendix S3.**
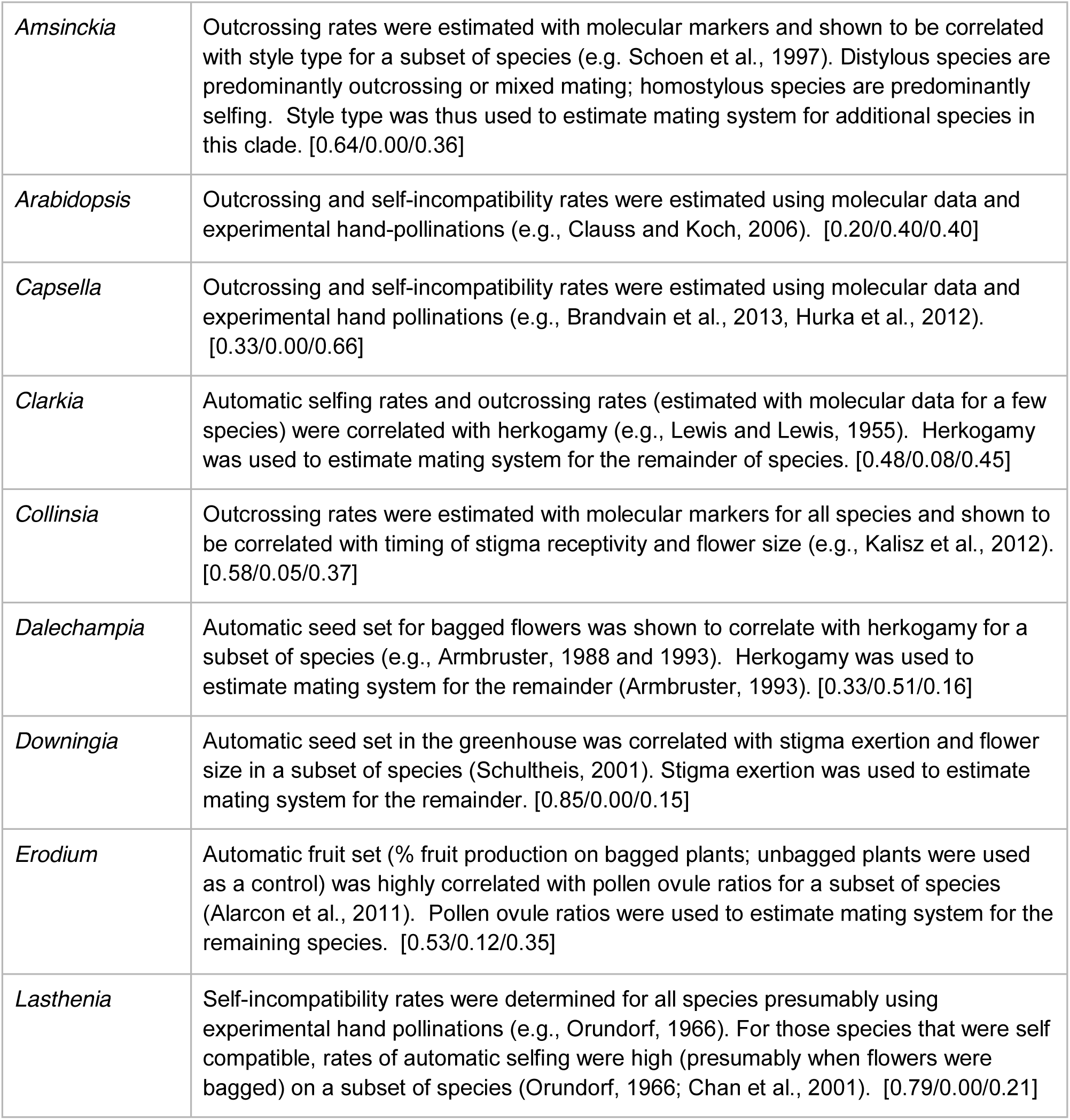
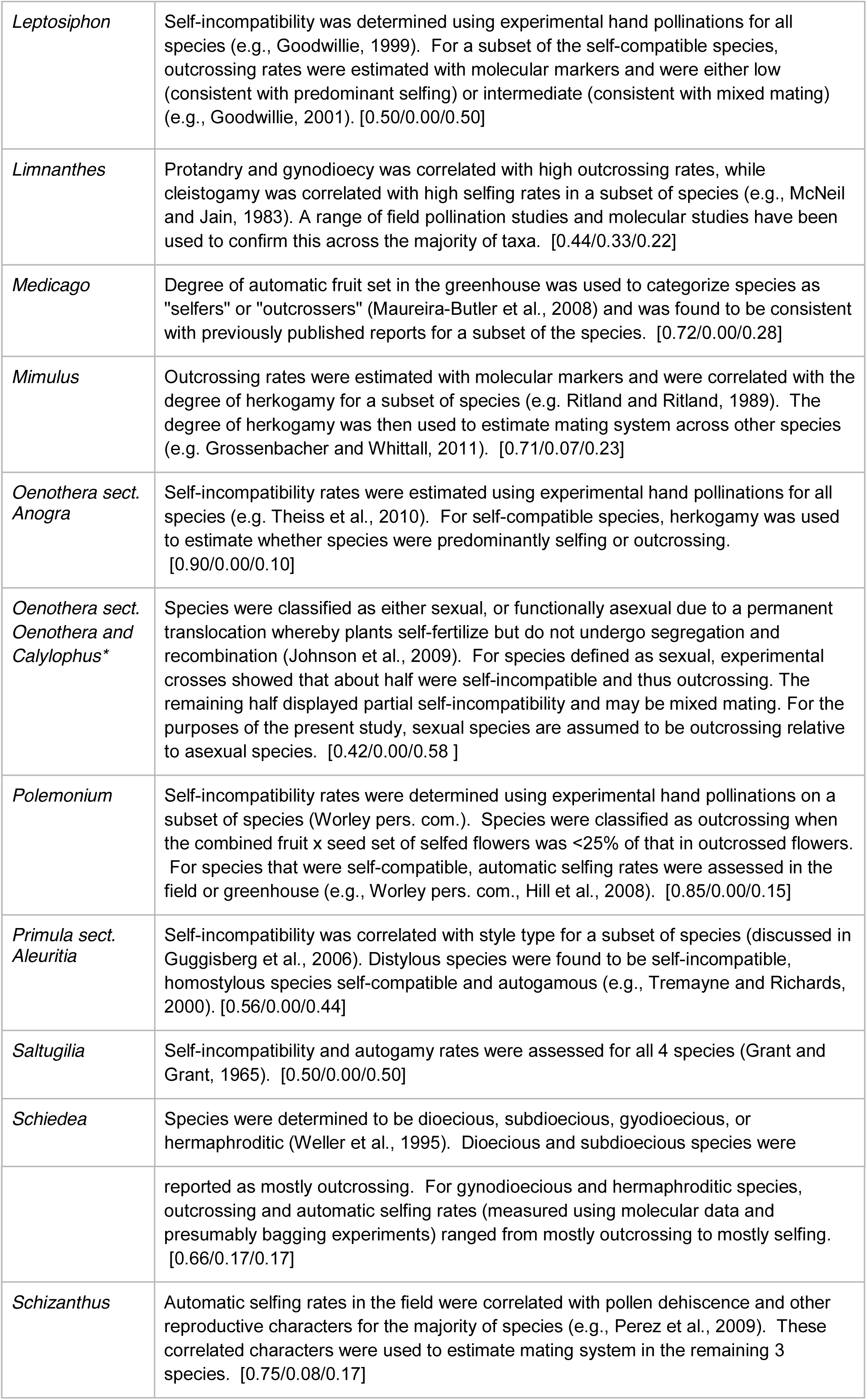
Description of how mating system was determined for each clade. The proportion of species assigned as outcrossers, variable maters, and selfers are included in brackets, [outcrosser/variable mater/selfer].

**Appendix S4.**
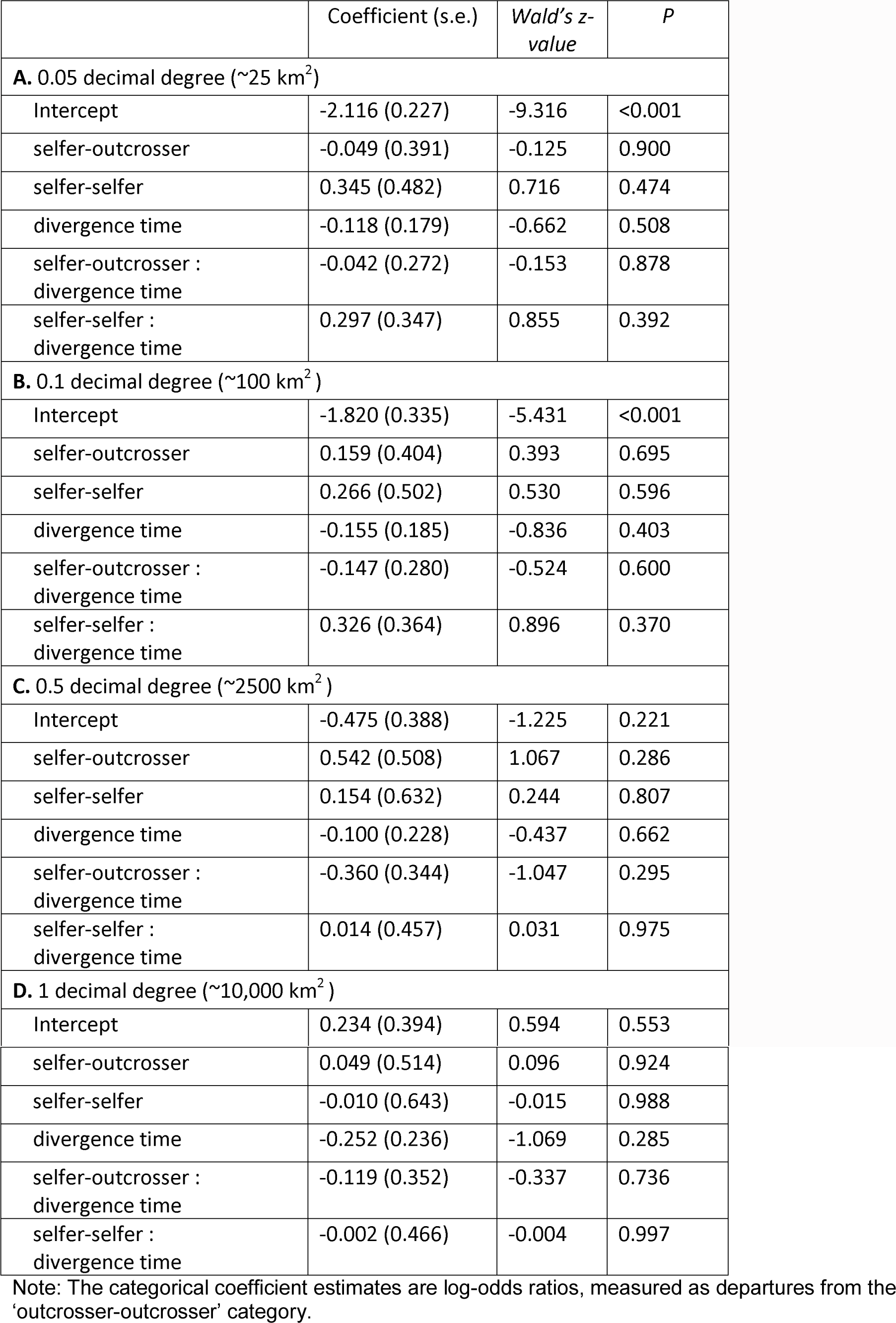
Results of beta regression models analyzing the effects of divergence time, ‘sister pair mating system’, and their interaction on co-occurrence, estimated at 822 four spatial scales (**A-D**).

